# Pitfalls of mapping functional and molecular human brain imaging data from separate cohorts

**DOI:** 10.1101/2025.04.22.650037

**Authors:** Patrick M. Fisher, Kristian Larsen, Pontus Plavén-Sigray, Gitte M. Knudsen, Brice Ozenne

## Abstract

It has become increasingly common to probe correlations between human brain imaging measures of receptor/protein binding and function using population-level brain maps drawn from independent cohorts, estimating correlations across regions. This strategy raises issues of interpretation that we highlight here with a multimodal brain imaging dataset and simulation studies. Twenty-four healthy participants completed neuroimaging with both [11C]Cimbi-36 positron emission tomography and magnetic resonance imaging scans to estimate receptor binding potential (BP) and cerebral blood flow (CBF), respectively, in 18 cortical/subcortical regions. Correlations between BP and CBF were estimated in three ways: 1) Pearson correlation across regions of mean regional BP and CBF (ρ_l_), to mimic studies using data from independent cohorts; 2) Pearson correlation between BP and CBF across participants in each region (ρ_2_); or 3) the correlation between BP and CBF across participants across all regions within a single linear mixed effects model (ρ_3_). We observed a significant positive correlation across regions (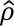_l_ = 0.672; p = 0.0023). Region-specific correlations across participants were substantively lower and not statistically significant (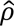_2_: mean = 0.140, range = -0.112 to 0.336; all p > 0.10), nor when estimated simultaneously within a linear mixed model (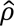_3_ = 0.138, p = 0.26). Our simulation study illustrated that regional differences in BP or CBF mean and variance can bias across regions correlations by 1000% or create a type-1 error of 100%. Our observations that both the estimated correlation and the statistical significance can differ greatly highlights that inferring across region correlation as evidence for correlation across participants is erroneous. Without validated methods that limit confounding and other biases, we discourage future studies from inferring across region correlation of population-level brain maps from independent cohorts in this manner.

## Introduction

Functional brain imaging tools, e.g., blood oxygen level dependent functional magnetic resonance imaging (BOLD fMRI), arterial spin labeling (ASL) MRI, electroencephalography, magnetoencephalography, and functional near-infrared spectroscopy, can be applied to assess individual task-related and task-independent activation and connectivity patterns across the brain under baseline conditions or following an intervention within healthy or clinical cohorts. Despite the wealth of brain-related information that can be gained from these tools, they are not without limitations. One such limitation is that it is not possible to directly derive the molecular signaling mechanisms (e.g., specific neurotransmitters or receptors) that contribute to an observed activation pattern or correlation structure. This information can be highly relevant for many reasons, e.g., knowledge of critical receptor pathways can guide a pharmacological intervention study targeting that receptor. Molecular brain imaging tools, e.g., positron emission tomography (PET) and single-photon emission compute tomography (SPECT) can illuminate the spatial distribution of specific neuromolecular processes, e.g., glucose metabolism, receptor binding or enzymatic activity. As such, acquiring both functional and molecular brain imaging data within a single cohort can implicate specific molecular mechanisms in shaping relevant brain function (Fisher & Hariri, 2013; Fisher et al., 2006; Sankar et al., 2023). Unfortunately, such a multimodal design is not feasible in many research settings. Molecular brain imaging is especially resource demanding, requiring a specialized infrastructure, i.e., cyclotron and radiochemistry laboratory, with costs that can be prohibitively expensive. Subsequently, researchers have sought to develop alternative strategies for mapping neuromolecular mechanisms onto brain activity and connectivity patterns.

The recent push to make more biomedical research data publicly available has contributed to the emergence of valuable resources such as the Allen Brain Atlas and other online repositories wherein one can obtain organ or brain-region specific levels of gene expression and/or protein concentrations. Similarly, human molecular imaging research groups are making publicly available more and more population-level brain-wide maps depicting, e.g., receptor or transporter levels (Beliveau et al., 2017; Johansen et al., 2024; Nørgaard et al., 2021; Savli et al., 2012). Subsequently, there has been a rapid expansion in studies leveraging these publicly available brain atlases, typically in conjunction with functional brain maps, to establish evidence of molecular mechanisms underlying aspects of brain function (see Supplementary Table 1 for non-exhaustive list of >40 such studies). In almost all cases, these association studies cannot be performed at the participant level because the two brain maps are derived from independent cohorts. Instead, a typical strategy is to average the functional and molecular brain maps across the respective participants for a common set of regions. A statistical analysis is then made *across regions* where a statistically significant correlation is inferred as evidence that the relevant molecular mechanism and functional brain signal are meaningfully related to one another in a manner that applies to broader populations, e.g., healthy individuals or specific diagnostics groups. Likewise, vice versa, the absence of evidence for an association between the two measures implicitly suggests a limited or negligible relation between the molecular and functional constructs. Concerns about an inflated type-1 error rate of this method have been noted previously and strategies aimed at addressing this limitation have been described, e.g., spatial permutation framework (Alexander-Bloch et al., 2018). However, there are reasons to believe that this interpretation of between-cohort correlations is fundamentally problematic and that such analyses do not inform about relations that are generalizable across individuals, irrespective of statistical significance correction strategies. Here we highlight these pitfalls through a multimodal human brain imaging dataset including serotonin 2A receptor (5-HT2AR) binding with [11C]Cimbi-36 PET and cerebral blood flow with ASL MRI and an associated simulation study. We also elaborate on the statistical principles underlying why this inference is problematic.

## Methods

### Participants

Here we report data from 24 healthy volunteers (10 females; age, mean ± sd [range] = 32.5 ± 7.9 [24-58] years) who participated in a broader study related to neural effects of serotonin 2A receptor (5-HT2AR) signaling (NCT03289949). Participants were generally healthy and without current or previous personal or family history of psychiatric or neurological disease. Detailed inclusion/exclusion criteria have been reported previously (Madsen et al., 2021). Measures and related information was extracted from the Cimbi database following an approved database application (https://nru.dk/index.php/allcategories/category/224-cimbi) (Knudsen et al., 2016). Prior to inclusion, participants received written and verbal descriptions of the study protocol; all participants provided written consent prior to study participation. The overall study was approved by the ethics committee for the Capital Region of Copenhagen (journal identifier: H-16028698, amendments: 56023, 56967, 57974, 59673, 60437, 62255) and Danish Medicines Agency (EudraCT identifier: 2016-004000-61, amendments: 2017014166, 2017082837, 2018023295). Cerebral blood flow and 5-HT2AR non-displaceable binding potential (BPND) estimates used in this article have been reported in previous, unrelated publications (Larsen et al., 2025; Søndergaard et al., 2022; Spies et al., 2020).

### Magnetic resonance imaging

The acquisition and preprocessing methods are described below and the same as described previously (Larsen et al., 2025). In brief, participants were scanned on one of two 3T Siemens Magnetom Prisma MRI scanners (Siemens AG, Erlangen, Germany) at Rigshospitalet, Copenhagen (MR1 and MR2). Data were drawn from two scanners because MR1 became unavailable during data collection. Fifteen participants completed scan sessions on MR1 and nine participants on MR2. All images were visually inspected for quality using the FSL image viewer, FSLeyes (Jenkinson, Beckmann, Behrens, Woolrich, & Smith, 2012).

### MR1 acquisition parameters

Imaging data were acquired using a 64-channel head/neck coil. We acquired a high-resolution, T1-weighted 3D MPRAGE structural image (inversion time = 900ms, echo time = 2.58ms, repetition time = 1900ms, flip angle = 9°, in-plane matrix = 256×256, in-plane resolution = 0.9mm×0.9mm, number of slices: 224, slice thickness = 0.9mm, no gap). We acquired a pseudo-continuous arterial spin labeling (pcASL) sequence to estimate CBF using a 2D echo-planar imaging (EPI) gradient spin-echo sequence (scan time = 332s, echo time = 12ms, repetition time = 4000ms, flip angle = 90°, label duration: 1508ms, single post-labeling delay = 1500ms, in-plane matrix = 64×64, in-plane resolution = 3mm×3mm, number of slices = 20, slice thickness = 5mm no gap, number of control/label pairs = 40, slice readout duration = 35ms). To calibrate the pcASL signal, an M0 scan was acquired using the same imaging parameters except for a repetition time of 10000ms. We acquired a gradient echo sequence (GRE) to estimate an underlying field map to correct for geometric distortions in the pcASL images (echo time 1 = 4.92ms, echo time 2 = 7.38ms, repetition time = 400ms).

### MR2 acquisition parameters

Imaging data were acquired using a 32-channel head coil. We acquired a high-resolution, T1-weighted 3D MPRAGE structural image (inversion time = 920ms, echo time = 2.41ms, repetition time = 1810ms, flip angle = 9°, in-plane matrix = 288×288, in-plane resolution = 0.8×0.8mm, slice thickness = 0.8mm, number of slices = 224, no gap). We acquired pcASL with a five post-label delay 3D turbo gradient spin echo sequence (scan time = 431s, 12 label/control pairs, native voxel size: 2.5×2.5×3mm3, image matrix = 96×96×40, label duration = 1508ms, post-label delays = [500, 500, 1000, 1000, 1500, 1500, 2000, 2000, 2000, 2500, 2500, 2500]ms, echo time = 3.78ms, repetition time = 4100ms, flip angle: 120°, two background suppression pulses optimised for each PLD). The acquisition included an M0 scan with which to calibrate the pcASL signal without any background suppression. Spin-echo echo-planar acquisitions with opposite phase-encoded blips were acquired to correct for geometric distortions using *topup* implemented in FSL (FMRIB software library, www.fmrib.ox.ac.uk) (Andersson, Skare, & Ashburner, 2003; Smith et al., 2004).

### Quantification of cerebral blood flow

### MR1 processing

The pcASL data underwent intramodal registration, dividing label and control images (40 pairs) before realigning them. The pcASL data were then unwarped to correct for B0 field distortions. Realignment and distortion correction was performed using SPM12 (https://Error**! Hyperlink reference not valid.** The high-resolution T1-weighted structural image was coregistered with the realigned and motion corrected pcASL data. We corrected for the interleaved slice acquisitions in the 2D EPI protocol using a delay in slice time acquisition of 35ms.

### MR2 preprocessing

For each PLD, a label/control difference image was realigned and the high-resolution T1-weighted structural image was co-registered with the M0 image. Co-registration with the pcASL images like we did for the MR1 data was not possible without introducing transformation artefacts.

All pcASL images from both scanners were quantified to absolute CBF values using BASIL in FSL (FMRIB software library, www.fmrib.ox.ac.uk), which estimates absolute CBF (ml/100g/min) using a single, well-mixed tissue compartment model (Buxton et al., 1998; Chappell, Groves, Whitcher, & Woolrich, 2009). We used standard settings for the quantitative modelling. Following segmentation of the high-resolution T1-weighted structural image in SPM12 to produce gray-matter, white-matter, and cerebrospinal fluid probability maps, mean gray-matter-probability-weighted regional CBF estimates were computed (see below for region definition).

### Positron emission tomography

[11C]Cimbi-36 was synthesized as previously reported (Ettrup et al., 2014). All PET scans were acquired using a Siemens ECAT High Resolution Research Tomograph (HRRT, CTI/Siemens, Knoxville, TN) in 3D acquisition mode and with an in-plane resolution of <2 mm. After securing the participant’s head in a head holder with cushions to limit motion, scans were performed following a bolus radioligand administration, using an established protocol: a 6-min transmission scan was followed by 120-min (frames: 6 × 10s, 6 × 20s, 6 × 60s, 8 × 120s, and 19 × 300s) of dynamic PET scanning, which commenced with [11C]Cimbi-36 injection (Ettrup et al., 2014, 2016). Scans were reconstructed using a 3D-OSEM-PSF algorithm (Hong et al., 2007; Sureau et al., 2008).

### Positron emission tomography processing

PET data were processed using Pvelab (Svarer et al., 2005). We applied the AIR algorithm to correct for motion during PET scans, wherein each PET scan was first smoothed with a 10 mm within-frame Gaussian filter prior to estimation of motion (Woods, Cherry, & Mazziotta, 1992). Motion-correction was applied to unfiltered PET frames. The high-resolution T1-weighted structural MRI scan was co-registered into PET space and segmented into gray-matter, white-matter, and cerebrospinal fluid probability maps using SPM12 (https://Error! **Hyperlink reference not valid.** Individual regions were defined in subject-space using Pvelab and included: middle/inferior frontal gyrus, superior frontal gyrus, and superior temporal gyrus as well as occipital cortex, orbitofrontal cortex, parietal cortex, anterior cingulate cortex, insula cortex, posterior cingulate cortex, ventrolateral prefrontal cortex, dorsolateral prefrontal cortex, sensory-motor cortex (i.e., pre/post-central gyri), amygdala, hippocampus, entorhinal cortex, thalamus, putamen, and caudate. A mean time series was computed for each grey-matter masked region to quantify [11C]Cimbi-36 non-displaceable binding potential (BPND) using the simplified reference tissue model (SRTM) with cerebellum as the reference region (Ettrup et al., 2014).

Subject-specific CBF images were co-registered to PET images using CBF-aligned M0 images and PET-aligned high-resolution T1-weighted structural images. Gray-matter-weighted regional CBF values were calculated within the 18 Pvelab-defined, subject-specific regions.

### On the Pearson correlation coefficient

The Pearson correlation coefficient quantifies the association between two variables based on a set of paired measurements (X, Y). The associated statistical test traditionally assumes that each pair of measurements is independent of each other (and identically distributed). This is a reasonable assumption across participants. For example, if we know the average [11C]Cimbi-36 BPND value for region A, based on a reference atlas (Beliveau et al., 2017), the region A BPND value of one participant provides no additional information about the region A BPND value for any other participants. However, this assumption is dubious across brain regions. For example, knowing that Participant 1 has a higher than average region A [11C]Cimbi-36 BPND value may suggest a similarly higher than average region B BPND value (Erritzoe et al., 2010). This between-region correlation is precisely why repeated-measures statistical models are commonly applied to brain imaging analyses (e.g., repeated measures ANOVA, linear mixed models, etc.). Therefore, with a two-level data structure (regional measurements nested into participants), the ordering of operations matters, i.e., the following two procedures are not equivalent: 1) first averaging over participants and then estimating the correlation across regions, which is similar to relating two data types from separate cohorts, or 2) estimating the correlation across participants and averaging those correlations over regions. The former, which relates two data types from separate cohorts, assumes independence across regions whereas the latter, which relates two data types from a single cohort, assumes independence across participants.

This two-level structure also leads to ambiguity about the definition of the correlation. Indeed, decomposing the residual variability into measurement noise and between-subject heterogeneity, one can distinguish between the *conditional* correlation (measurement noise only) and the *marginal* correlation (measurement and subject noise). Typically, studies aim to estimate the marginal correlation, e.g., estimating the correlation between 5-HT2AR binding and CBF in a region over a large, possibly heterogeneous, population. In contrast, with the former, one would focus on a homogenous sub-group of the population such as young healthy adults with similar genetic state. In practice, researchers want to leverage measurements over multiple regions to make up for the limited number of participants that can be included due to financial and logistic constraints.

### Statistical analyses

All statistical analyses were performed in R (version 4.3.1) (R Core Team, 2023). We compared three different strategies for estimating the Pearson correlation coefficient between 5-HT2AR binding and CBF:

- *1) Across regions strategy*: compute mean regional BPND and CBF across all participants; estimate the correlation between BPND and CBF across regions (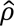_l_) This represents what is typically done when correlating data drawn from two separate cohorts.
- *2) Across participants strategy*: compute the correlation between the observed BPND and CBF across participants, for each region, separately 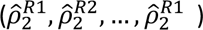. This represents a strategy for correlating two imaging measures drawn from a single cohort.
- 3) *Mixed model strategy*: the correlation, 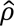_3_, between the observed BPND and CBF across participants for all regions is fit with a single linear mixed model where each participant has 36 measurements, i.e., 18 regions and two modalities (i.e., BPND and CBF). The model includes a mean and variance parameter specific to each region and modality, as well as four correlation parameters. See Supplementary Material A1 for additional model details. This strategy is closely related to Strategy 2 but estimates the average marginal correlation instead of region-specific marginal correlations.

Both Strategies 1) and 2) estimate correlation using the standard implementation of the Pearson correlation coefficient (i.e., *cor.test* from the *stats* package: https://www.rdocumentation.org/packages/stats/versions/3.6.2). Strategy 3 corresponds to a specific residual covariance pattern but can also be fitted as a random effect model when the correlation is positive (see Supplementary Material A1).

Notably, studies combining data from separate cohorts can employ only Strategy 1. Here we leverage that the two imaging data types were acquired in the same cohort to evaluate and compare results from all three strategies. Statistical significance was defined as p < 0.05, all reported p-values are not adjusted for multiple comparisons.

### Simulation study

We performed a simulation study to further assess the effect of confounding bias and misinterpretation of inferring across region correlation as evidence for across participant correlation. We assessed the bias of the estimated correlation, coverage of the 95% confidence intervals, and type-1 error control and statistical power vis-à-vis p < 0.05 of Strategy 1-3 under five scenarios (A-E) for regional difference in average signal and signal variability (see Table 1 for simulation details). BPND and CBF values were simulated using a multivariate normal distribution with mean, variance, and correlation values estimated based on the observed data, except for the correlation between BPND and CBF in Scenario A-D which was either set to 0 or to a value larger than the observed. Scenario A and C and E simulates non-zero correlation between BPND and CBF, whereas Scenario B and D simulates no correlation between BPND and CBF. Correlations for Scenario A and C were selected arbitrarily (marginal/conditional correlation = 0.250/0.500), whereas Scenario E correlation reflects the observed data (marginal/conditional correlation = 0.135/0.057).

**Table 1.**
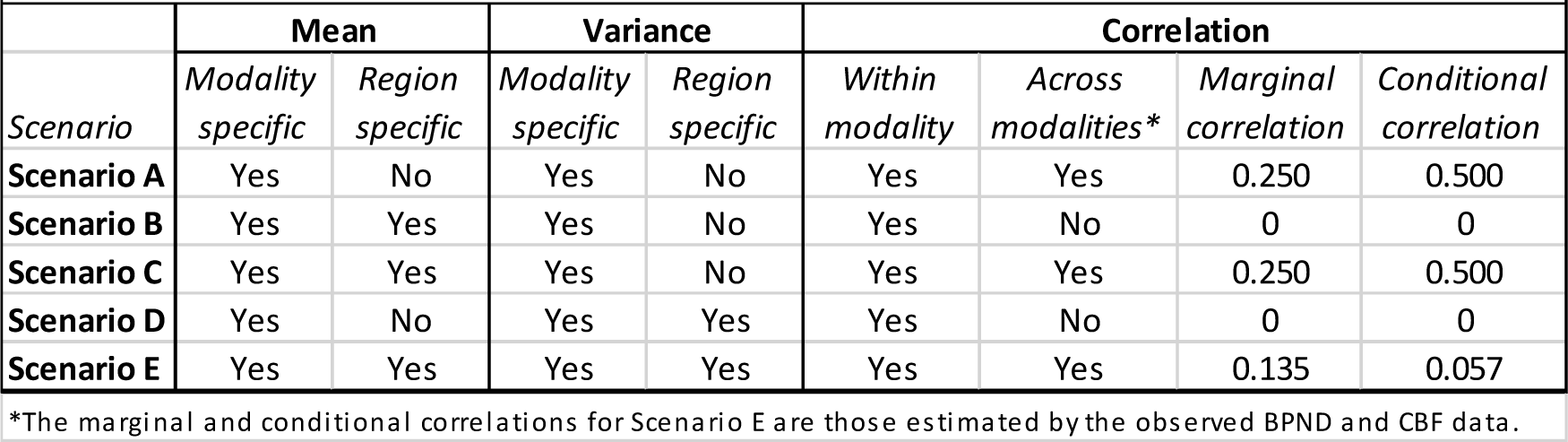
Simulation scenarios.

Each of the above scenarios was simulated 10000 times using the same structure as our real data, i.e., 18 regions, two measurements (BPND and CBF) and either 1) 24 participants (as in our study) or 2) 1000 participants. For Strategy 2, the average of the region-specific correlations was used and its uncertainty was quantified assuming that the Fisher transformed region-specific correlations are jointly normally distributed with variance 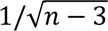 and correlation estimated via the influence function described in Equation 3 of (Devlin, Gnanadesikan, & Kettenring, 1975). For Strategy 3, the mixed model included a modality-and region-dependent mean but only a modality-dependent variance (to save computation time), meaning that Strategy 3 was mis-specified under Scenario D and E. No missing data was considered. A more elaborate description of the data generating mechanism for Simulation Scenario A-E can be found in Supplementary Material A2.

Code used to fit statistical models estimating the correlation coefficients and simulate data can be found here (https://github.com/bozenne/article-mapping-fMRI-PET).

## Results

### Across regions strategy (Strategy 1)

Employing the strategy applied when combining data from separate cohorts, we observed a statistically significant positive correlation between 5-HT2AR binding and CBF (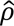_l_[95% CI] = 0.672 [0.298, 0.867]; p = 0.0023; Figure 1).

**Figure 1:**
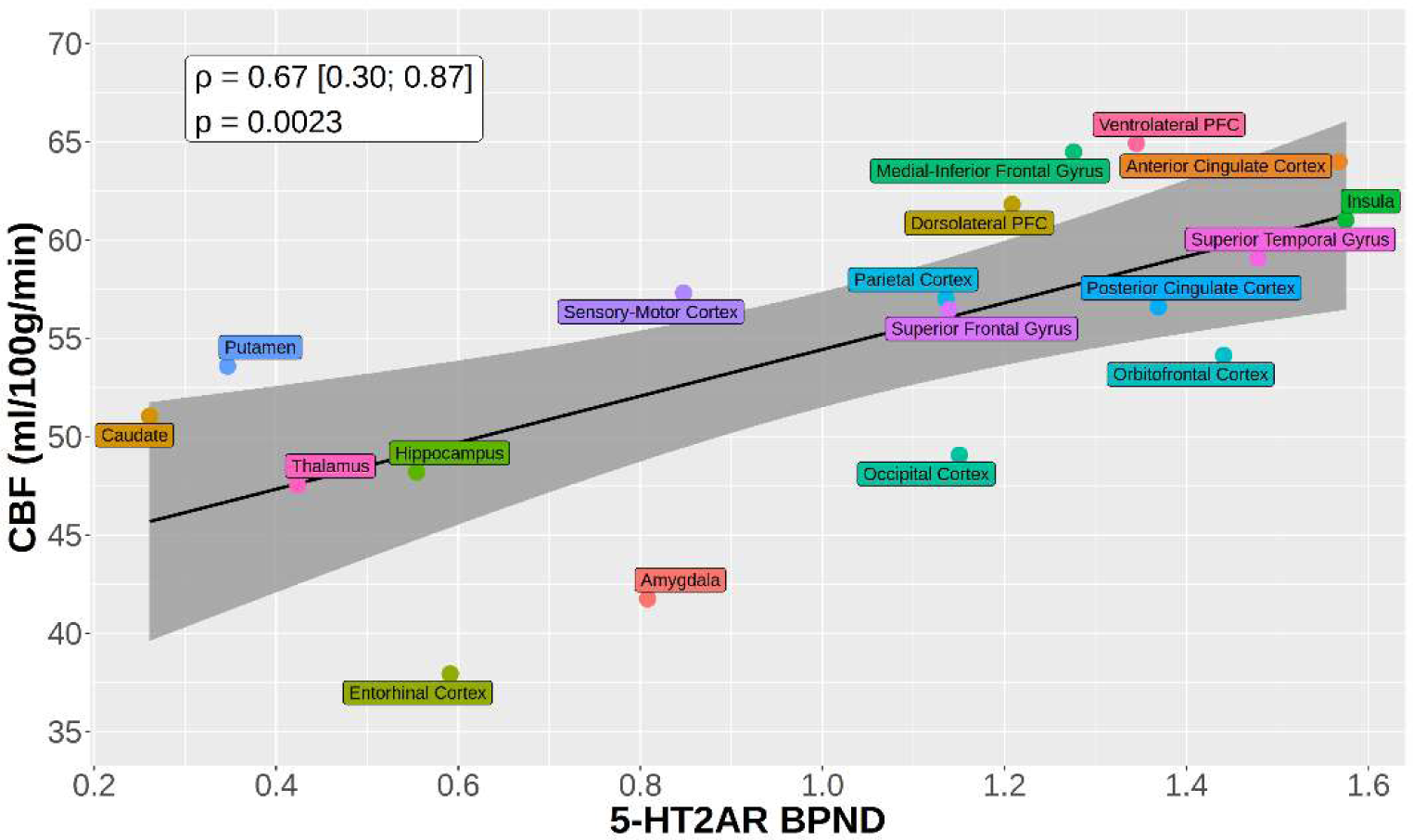
Across regions correlation between CBF and 5-HT2AR BPND. For each region, average CBF or 5-HT2AR BPND value was calculated across participants. The p-value and confidence interval for the Pearson correlation coefficient are estimated without consideration of possible correlation between the regional values from the same modality.

### Across participants strategy (Strategy 2) and mixed model strategy (Strategy 3)

Leveraging that the 5-HT2AR binding and CBF data were acquired in the same cohort, we evaluated the correlation across participants for each region, separately. The estimated correlations between 5-HT2AR binding and CBF ranged from -0.112 to 0.336 (average 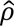_2_ = 0.140, Figure 2), none were statistically significant (Table 2). A similar result was obtained when using a linear mixed model to estimate the correlation between CBF and BPND across participants with region as a repeated measure (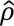_3_ = 0.138 [-0.100, 0.362]; p=0.26).

**Figure 2:**
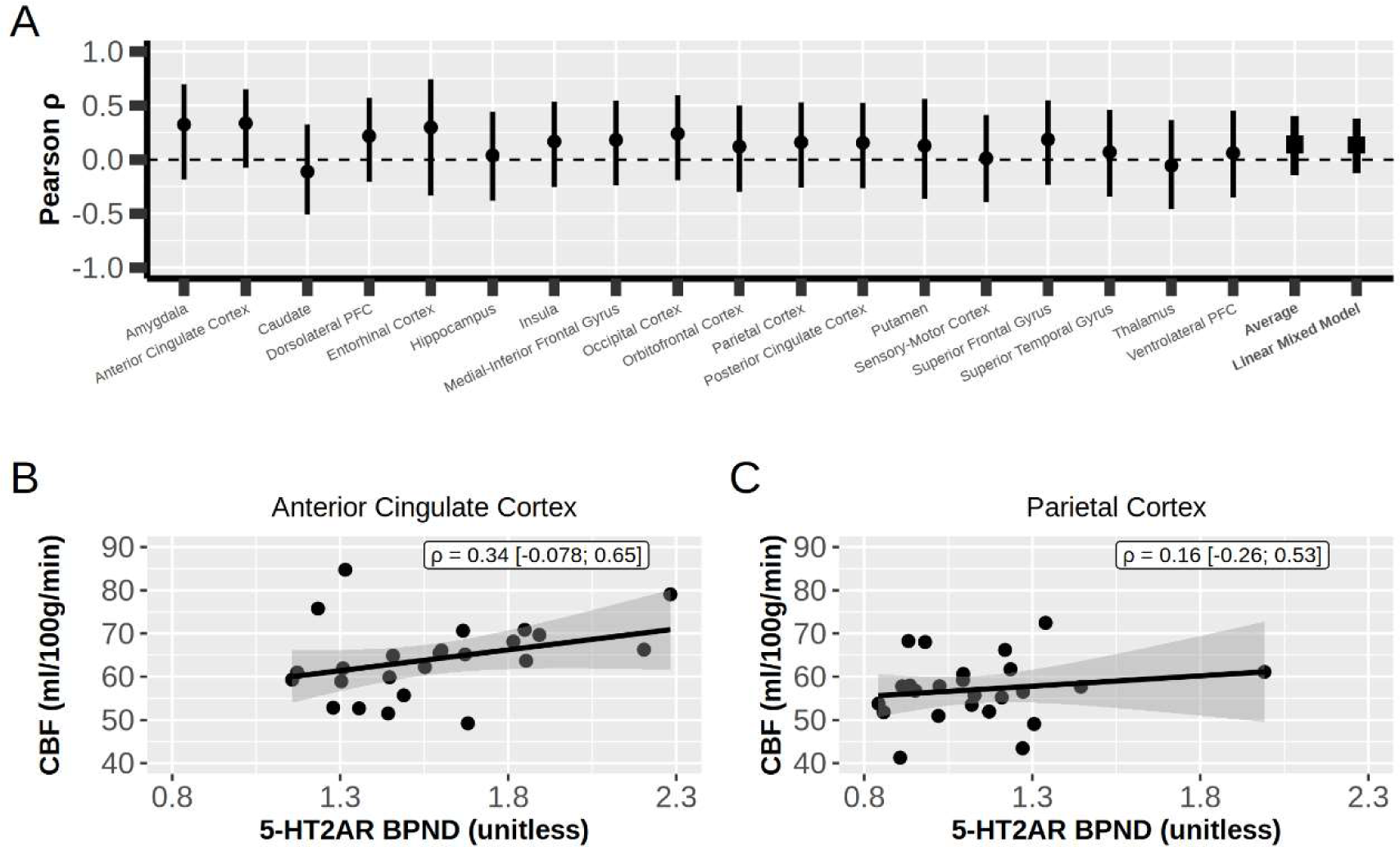
Across participants correlations between CBF and 5-HT2AR BPND. A) The Pearson correlation coefficient across participants for each region. Points and error bars denote correlation coefficient and 95% confidence interval, respectively. B) and C) Regional correlations between 5-HT2AR binding (BPND) and cerebral blood flow (CBF) for anterior cingulate and parietal cortex, respectively. Estimated Pearson correlation and 95% confidence interval in text box.

**Table 2.** Regional correlation coefficients between [11C]Cimbi-36 BPND and CBF.

| Region | Correlation | Lower | Upper | P-value |
| --- | --- | --- | --- | --- |
| Amygdala | 0.325 | -0.185 | 0.697 | 0.20 |
| Anterior cingulate cortex | 0.336 | -0.078 | 0.651 | 0.11 |
| Caudate | -0.112 | -0.51 | 0.325 | 0.62 |
| Dorsolateral prefrontal cortex | 0.217 | -0.204 | 0.571 | 0.31 |
| Entorhinal cortex | 0.297 | -0.333 | 0.744 | 0.35 |
| Hippocampus | 0.039 | -0.379 | 0.444 | 0.86 |
| Insula | 0.167 | -0.254 | 0.534 | 0.44 |
| Medial-inferior frontal gyrus | 0.182 | -0.239 | 0.545 | 0.39 |
| Occipital cortex | 0.240 | -0.191 | 0.594 | 0.27 |
| Orbitofrontal cortex | 0.120 | -0.298 | 0.499 | 0.58 |
| Parietal cortex | 0.160 | -0.260 | 0.529 | 0.46 |
| Posterior cingulate cortex | 0.155 | 0.265 | 0.526 | 0.47 |
| Putamen | 0.128 | -0.360 | 0.561 | 0.61 |
| Sensory-Motor cortex | 0.012 | -0.393 | 0.413 | 0.96 |
| Superior frontal gyrus | 0.186 | -0.235 | 0.548 | 0.38 |
| Superior temporal gyrus | 0.069 | -0.344 | 0.460 | 0.75 |
| Thalamus | -0.055 | -0.457 | 0.365 | 0.80 |
| Ventrolateral prefrontal cortex | 0.060 | -0.352 | 0.453 | 0.78 |
| **Average** | **0.140** | **-0.143** | **0.403** | **0.33** |
| **Linear mixed model (marginal)** | **0.135** | **-0.125** | **0.379** | **0.31** |

### Simulation study

Table 3 summarizes the simulation study under each scenario, including a description of the bias in estimated correlation coefficient, coverage (i.e., fraction of times simulated 95% CI includes the true correlation), type-1 error control (i.e., fraction of times simulated p < 0.05 where null hypothesis is true, relevant for Scenario B and D) and statistical power (i.e., fraction of times simulated p < 0.05 where null hypothesis is false, relevant for Scenario A, C and E). In Scenario A, where there was a true correlation between CBF and BPND and no regional differences, all strategies provided a nearly unbiased estimator (i.e., all biases < 2% different from ground truth). Where underlying regional differences in BPND and CBF are introduced (Scenarios B and C), Strategy 1 (i.e., strategy mimicking data drawn from separate cohorts) shows substantial bias, essentially no type-1 error control in Scenario B (i.e., 100% rejection rate where the null hypothesis is true). Conversely, Strategy 2 and 3 retain low bias (Scenario 3: ≤ 2% different from ground truth), type-1 error control close to nominal level (Scenario 2: 4.8%-5.7%), and statistical power that increases when the sample size is increased from N = 24 to N = 1000 participants (Scenario C). Notably, the 100% coverage of Strategy 1 in Scenario C was due to wide confidence intervals relative to the bias (width = 0.570, bias = 0.170). Scenario D, which simulated data in the absence of a correlation between BPND and CBF and regional differences in variance but not in mean, showed good coverage and type-1 error control for all three strategies. Scenario E, which simulated data with correlations between BPND and CBF and regional differences in mean and variance, showed a substantial bias for Strategy 1 (>1000%) and no coverage, i.e., the correlation coefficient was never well estimated. The model misspecification of Strategy 3 in Scenario E resulted in a negative bias (∼10%) and some under-coverage (90% instead of 95%) for N = 1000 participants. Notably, statistical power increases with sample size (N = 24 to 1000) for Strategy 2 and 3 under all relevant scenarios (A, C, and E), but not Strategy 1. This is because the “sample size” for Strategy 1 is how many regions are defined, not how many participants are sampled. See Supplementary Material A3 for a visual representation of simulation scenarios.

**Table 3.**
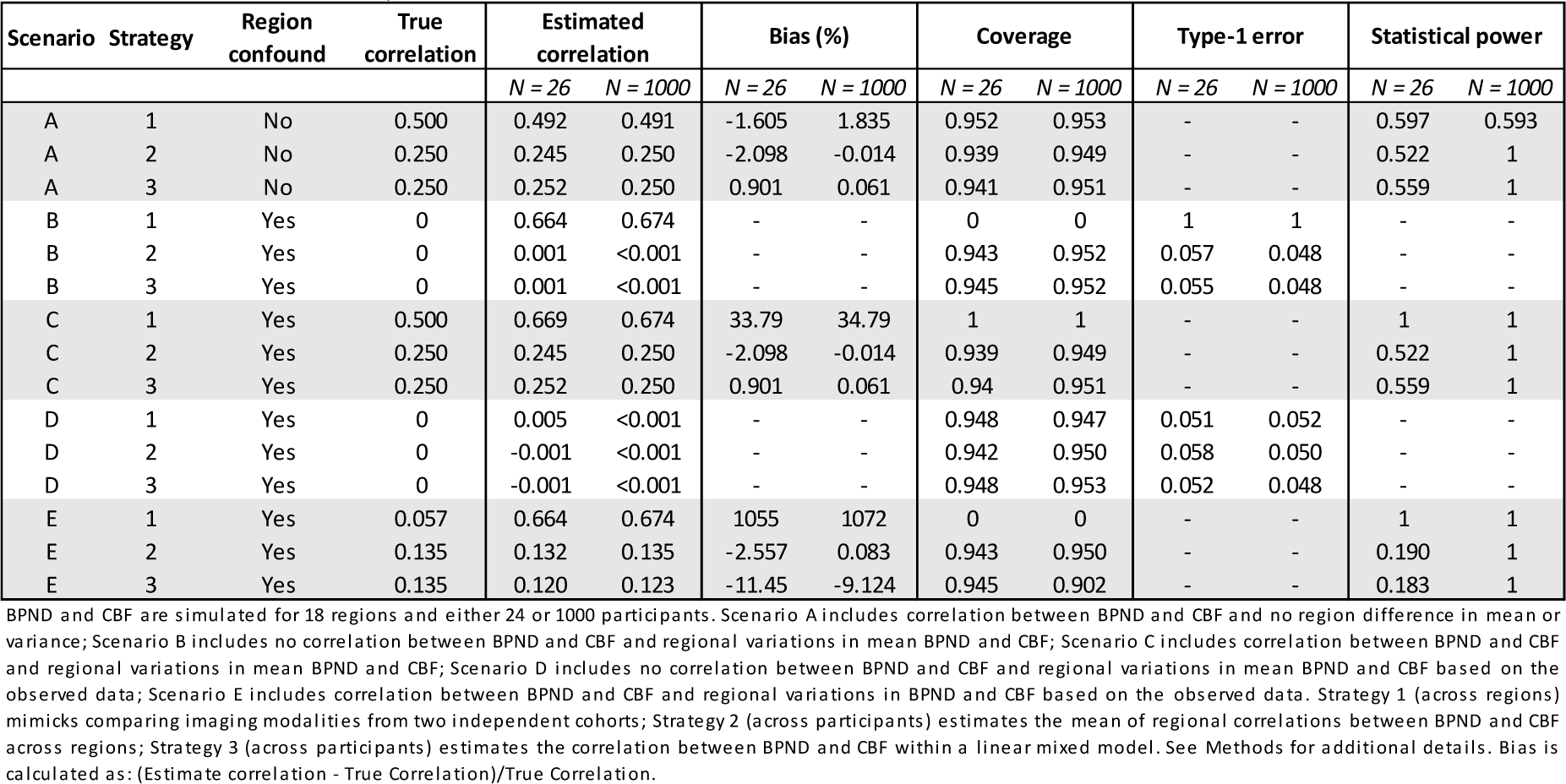
Results from simulation study scenarios.

| Scenario | Strategy | Region confound | True correlation | Estimated correlation |  | Bias (%) |  | Coverage |  | Type-1 error |  | Statistical power |  |
| --- | --- | --- | --- | --- | --- | --- | --- | --- | --- | --- | --- | --- | --- |
|  |  |  |  | N = 26 | N = 1000 | N = 26 | N = 1000 | N = 26 | N = 1000 | N = 26 | N = 1000 | N = 26 | N = 1000 |
| A | 1 | No | 0.500 | 0.492 | 0.491 | -1.605 | 1.835 | 0.952 | 0.953 | - | - | 0.597 | 0.593 |
| A | 2 | No | 0.250 | 0.245 | 0.250 | -2.098 | -0.014 | 0.939 | 0.949 | - | - | 0.522 | 1 |
| A | 3 | No | 0.250 | 0.252 | 0.250 | 0.901 | 0.061 | 0.941 | 0.951 | - | - | 0.559 | 1 |
| B | 1 | Yes | 0 | 0.664 | 0.674 | - | - | 0 | 0 | 1 | 1 | - | - |
| B | 2 | Yes | 0 | 0.001 | <0.001 | - | - | 0.943 | 0.952 | 0.057 | 0.048 | - | - |
| B | 3 | Yes | 0 | 0.001 | <0.001 | - | - | 0.945 | 0.952 | 0.055 | 0.048 | - | - |
| C | 1 | Yes | 0.500 | 0.669 | 0.674 | 33.79 | 34.79 | 1 | 1 | - | - | 1 | 1 |
| C | 2 | Yes | 0.250 | 0.245 | 0.250 | -2.098 | -0.014 | 0.939 | 0.949 | - | - | 0.522 | 1 |
| C | 3 | Yes | 0.250 | 0.252 | 0.250 | 0.901 | 0.061 | 0.94 | 0.951 | - | - | 0.559 | 1 |
| D | 1 | Yes | 0 | 0.005 | <0.001 | - | - | 0.948 | 0.947 | 0.051 | 0.052 | - | - |
| D | 2 | Yes | 0 | -0.001 | <0.001 | - | - | 0.942 | 0.950 | 0.058 | 0.050 | - | - |
| D | 3 | Yes | 0 | -0.001 | <0.001 | - | - | 0.948 | 0.953 | 0.052 | 0.048 | - | - |
| E | 1 | Yes | 0.057 | 0.664 | 0.674 | 1055 | 1072 | 0 | 0 | - | - | 1 | 1 |
| E | 2 | Yes | 0.135 | 0.132 | 0.135 | -2.557 | 0.083 | 0.943 | 0.950 | - | - | 0.190 | 1 |
| E | 3 | Yes | 0.135 | 0.120 | 0.123 | -11.45 | -9.124 | 0.945 | 0.902 | - | - | 0.183 | 1 |

BPND and CBF are simulated for 18 regions and either 24 or 1000 participants. Scenario A includes correlation between BPND and CBF and no region difference in mean or variance; Scenario B includes no correlation between BPND and CBF and regional variations in mean BPND and CBF; Scenario C includes correlation between BPND and CBF and regional variations in mean BPND and CBF; Scenario D includes no correlation between BPND and CBF and regional variations in mean BPND and CBF based on the observed data; Scenario E includes correlation between BPND and CBF and regional variations in BPND and CBF based on the observed data. Strategy 1 (across regions) mimicks comparing imaging modalities from two independent cohorts; Strategy 2 (across participants) estimates the mean of regional correlations between BPND and CBF across regions; Strategy 3 (across participants) estimates the correlation between BPND and CBF within a linear mixed model. See Methods for additional details. Bias is calculated as: (Estimate correlation - True Correlation)/True Correlation.

## Discussion

Recent studies have sought to exploit human brain imaging datasets derived from different cohorts to, e.g., map neuromolecular phenotypes onto brain function or connectivity (Supplementary Table 1). The results are then often interpreted as evidence for associations between these two measures that generalize to larger populations. Here we analyze a multimodal human brain imaging dataset to provide a counterfactual example of this inference to highlight the pitfalls of incorrectly interpreting correlations estimated across regions where imaging data are acquired from separate cohorts. We observed a statistically significant positive correlation between 5-HT2AR BPND, measured with [11C]Cimbi-36 PET, and CBF, measured with ASL MRI, when modeled across regions, representing an analysis of brain imaging measures acquired in separate cohorts. However, we did not observe evidence for a significant correlation between these two measures for any region, nor in a linear mixed model framework with region as a repeated measure, when modeled across participants, representing an analysis where both imaging data are acquired in the same cohort. Our simulation study highlighted that the across region correlation variably and sometimes substantially misestimates the true correlation and can show poor type-1 error control. Hence, the across region strategy is particularly vulnerable in situations such as where there are regional differences in mean signals, which is common for brain imaging measures. Although our observations do not demonstrate that *all* across-region correlations exist in the absence of an across-participant correlation, it emphasizes that these two types of correlations represent two distinct constructs that are not reliably related. As such, it is erroneous to conclude that an across-region correlation is indicative of an across-participant correlation, which raises concerns about the interpretation of a growing number of studies that integrate imaging data from two separate cohorts.

A core challenge with the across region analysis strategy can be understood by considering the underlying assumptions of the Pearson correlation coefficient, ρ. This can be effectively estimated based on *independent* replicates of two variables, X and Y, that are not confounded. The across-region strategy, e.g., Strategy 1, grossly violates this assumption in many (if not all) multimodal human brain imaging analyses. For instance, regional PET measurements are typically highly correlated within a participant (Erritzoe et al., 2010) and both PET and fMRI show substantial regional variations in population mean value (Figure 1). Our simulations show that the correlation estimated by Strategy 1 (across regions) can be very sensitive to regional mean differences and variably biased with respect to the correlation structure across participants. The 5-HT2AR and CBF data used here highlight this confound: there are regional differences in both 5-HT2AR and CBF and they have a similar regional rank order structure. Subsequently, for our observed dataset, we observed a significant correlation across regions (Strategy 1), whereas the regional correlations across participants (Strategy 2 and 3) were substantially lower and not statistically significant. Our simulation results suggest the breadth of circumstances where these biases affect estimated correlations. In Scenario B there is no underlying correlation between BPND and CBF, yet a strong correlation coefficient is ubiquitously estimated by the across regions strategy. Subsequently, the type-1 error rate, which should be 5%, is instead 100%. In Scenario C, where there is a true underlying correlation between BPND and CBF, the across regions strategy (Strategy 1) consistently misestimates this correlation. Although Strategy 1 correctly estimates a null effect in Scenario D, it has a bias of >1000% in Scenario E, where the regional 5-HT2AR binding and CBF differences are equivalent. The erratic bias of Strategy 1 across scenarios highlights that it is highly susceptible to error, irrespective of whether there is a true underlying correlation. Alternative strategies are not without problems, which is illustrated in Scenario E where the variance was incorrectly modeled by the linear mixed model. Specifying the variance and correlation via a linear mixed model can be challenging, especially with fewer participants compared to the number of regions. Nevertheless, a reasonable model structure showed some robustness to misspecification, particularly vis-à-vis the dramatic effect of ignoring confounding by region.

Notably, from our simulated data, we show that the across regions strategy estimates a high correlation coefficient in the presence of regionally variable estimates, irrespective of whether there is (Simulation Scenario C) or is not (Simulation Scenario B) a true underlying correlation between the two estimates. Essentially, this means that it is very difficult, or perhaps not even possible, to dissociate the correlation due to this confound and true correlation between measures across participants. This bias is particularly problematic vis-à-vis a commonly applied spatial permutation framework (Alexander-Bloch et al., 2018). This permutation-based approach adjusts only the estimate of statistical significance, whereas we show that the correlation itself is variably and sometimes substantially biased, limiting its interpretability, irrespective of the p-value. It is not currently clear to us how one can estimate this confound between two imaging measures without measuring both in a single cohort, which limits the application of datasets from different cohorts. Indeed, controlling for confounding typically requires individual measurement of the confounder, exposure, and the outcome. It worth remembering that a linear model regressing, e.g., BPND on CBF (or vice versa), estimates a slope parameter that is the Pearson correlation rescaled by the standard deviation of the outcome and the exposure, i.e., 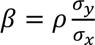, and the corresponding Wald test statistic can be shown to be a function of the Pearson correlation and the sample size, i.e., 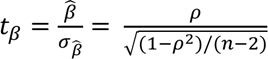. Put another way, concerns about confounding when fitting correlation coefficients also apply to fitting a linear regression.

Another perspective that highlights the limitation of exchanging across regions correlations for evidence of across participants correlations can be seen through the variance-covariance matrix that emerges from a linear mixed model fit when data are from a single cohort (see Supplementary Material A1). From this model we can estimate the correlation between BPND and CBF from the same region (*r_region_*). By contrast, the across region correlation is not directly estimated within the variance-covariance matrix, suggesting that it captures a correlation not straight-forwardly aligned with how correlation is typically understood by practitioners nor estimated when data are drawn from a single cohort.

An observation from the simulation study that highlights the conceptual peculiarity of the across regions correlation can be seen in the statistical power. In Simulation Scenario A and C, where a true correlation exists, the statistical power of the across regions strategy (Strategy 1) is unchanged increasing from N = 24 to 1000 participants. That the number of participants observed is essentially irrelevant to the uncertainty of the correlation estimate, and subsequently the p-value, speaks to how this does not reflect a correlation estimate that is generalizable to broader populations. Conversely, and similarly counter-intuitively, increasing the number of regions, i.e., parcellating the brain into smaller units, *would* increase the statistical power precisely because it is a correlation across regions.

Here we use an available multimodal brain imaging dataset to highlight the limitations of this across regions analysis strategy. The simulation study results reinforce that these pitfalls are not unique to the idiosyncratic features of the current empirical dataset: 1) although CBF and [11C]Cimbi-36 PET likely have their own susceptibility “fingerprint”, the underlying statistical problem generalizes to other measures, such as BOLD fMRI task activation and connectivity; 2) sample size, our simulation studies show that across regions correlations are counter-intuitively independent of participant sample size; 3) region set, in fact, increasing the number of regions will generally amplify the type-1 error, which increased to 100% with only 18 regions in our simulation study; and 4) statistical model, here we applied a simple linear regression across regions (Strategy 1), but the same inferential misinterpretation extends to multivariate estimations (e.g., canonical correlation analysis) of correlation/association.

One may argue that because multimodal studies are sometimes prohibitively expensive or impractical, estimating across regions correlations from separate cohorts is “better than nothing.” The absence of a mathematical link between across region and across participant correlations and the erratic divergence from known correlations that we observed across our empirical data and simulation studies suggests these results are unpredictably misleading and therefore not clearly useful in the absence of multimodal data.

In summary, we present a counterfactual example that challenges the increasingly published interpretation that an across region correlations between imaging datasets from separate cohorts constitutes evidence for a relation between the two imaging measures that generalizes to broader populations. In addition to misleading the assessment of molecular mechanisms underlying functional brain imaging phenotypes, misinterpretation of these across region correlations could facilitate, e.g., human pharmacological studies focusing on drug targets erroneously determined to be related to an imaging measure or clinical phenotype. We thus recommend researchers and associated stakeholders be mindful of these pitfalls.

## Data and Code Availability

Code for analyses can be found here: https://github.com/bozenne/article-mapping-fMRI-PET. Individual participant data can be made available with an appropriate data sharing agreement; please contact the corresponding author for more information.

## Author Contributions

PF: Conceptualization, Methodology, Software, Validation, Formal analysis, Investigation, Data Curation, Writing – Original Draft, Writing – Review & Editing, Supervision

KL: Investigation, Data Curation, Writing – Review & Editing, Visualization, Supervision

PPS: Writing – Review & Editing

GMK: Writing – Review & Editing, Funding acquisition

BO: Methodology, Software, Validation, Formal analysis, Writing – Original Draft, Writing – Review & Editing, Visualization, Supervision

## Acknowledgements

We gratefully acknowledge the work of MRI assistants for their assistance with data collection. We thank Lone Freyr, Gerda Thomsen, Svitlana Olsen, Peter Jensen, and Dorthe Givard for technical/administrative assistance. We would like to thank the Department of Radiology at Rigshospitalet for MR1 scanner access. We would like to thank the Kirsten & Freddy Johansen (KFJ) Foundation for funding the MR2 scanner.

## SUPPLEMENTARY MATERIAL

**Supplementary Table 1.** Example studies that evaluate across region correlations or estimations with spatial brain maps from two separate cohorts for the purpose of inferring generalizable relations between functional and molecular maps. Additional citation information can be found in Supplementary References.

**Supplementary Table 1.**
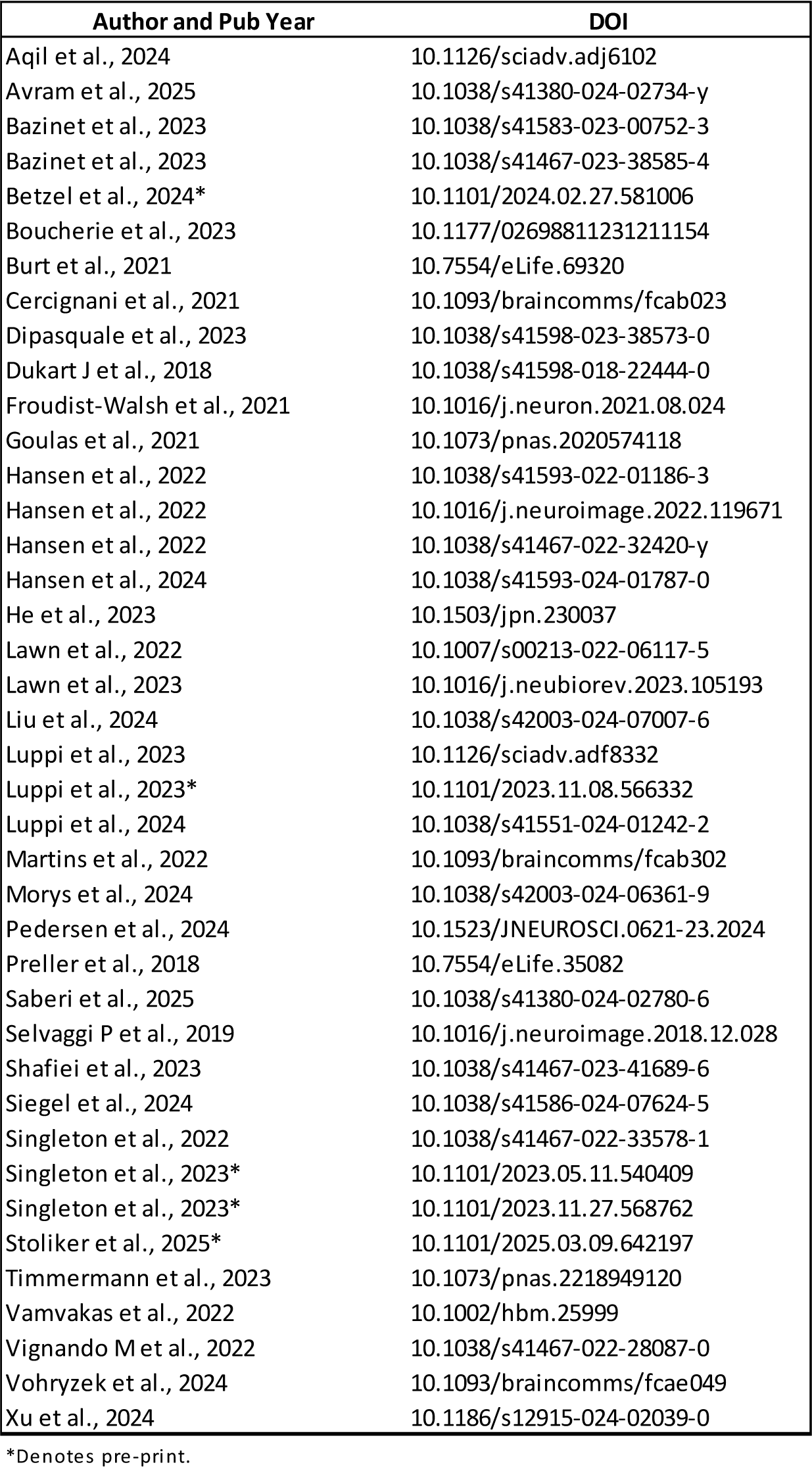
Studies estimating correlations or associations between pairs of spatial brain maps from different cohorts.

### Supplementary Material A1

Specification of the linear mixed model

The proposed linear mixed model (Strategy 3) uses four correlations parameters to model the within and between modality correlation:

- *r*_BPND_: correlation between BPND measurements from different regions
- *r*_CBF_: correlation between CBF measurements from different regions
- *r*_region_: correlation between a BPND and a CBF measurement from the same region
- *r*_id_: correlation between a BPND and a CBF measurement from different regions

The resulting correlation matrix is shown below, considering three example regions (thalamus, putamen, caudate):

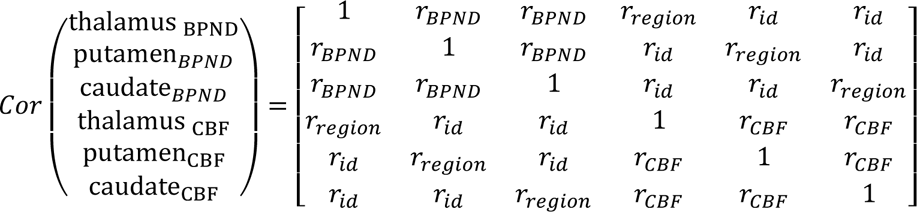

Based on the Restricted Maximum Likelihood estimates of the correlation parameters, the correlation of interest was estimated as 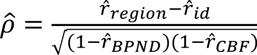. This formula has been derived from an alternative, and more restrictive, formulation of the model using random effects, where for region *r*:

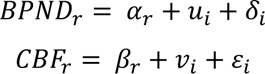

α and β; denote the region-specific intercepts, *u* and *v* denote the participant- and modality-specific random effects, and δ and ε, the residuals. Random effects and residuals are modeled to be independent from each other:

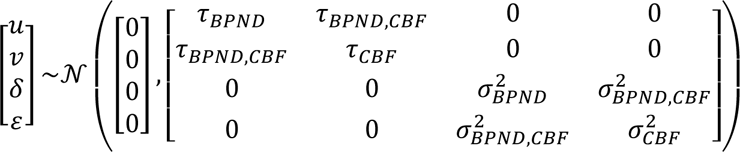

It can then be shown when *r*_BPND_ > 0, *r*_CBF_ > 0, *r*_region_ > *r*_id_ > 0 and assuming that the variance is not region dependent, but may be modality dependent, the proposed mixed model and the random effect models are equivalent and 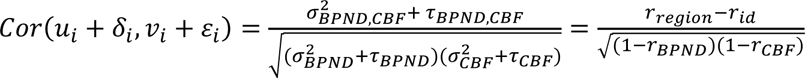

### Supplementary Material A2

Simulation study data generating mechanism

The data were simulated considering 18 regions and using a multivariate normal distribution parametrized according to the estimated parameters of a linear mixed model with modality and region dependent mean, modality dependent variance (Scenario A, B, C) or modality and region dependent variance (Scenario D, E), and the correlation structure described in Supplementary Material A1. This leads to the following values:

- Mean: Scenario A and D: mean BPND of 1.027 and mean fMRI of 54.772; Scenario B, C, E: region dependent means, BPND: 0.242 to 1.575; and CBF: 37.933 to 64.914.
- Standard deviation: Scenario A, B; C: standard deviation BPND of 0.255 and standard deviation CBF of 9.270; Scenario D, E: region dependent standard deviations, BPND: 0.105 to 1.399; and CBF: 6.355 to 16.998.
- Within-modality correlation: Scenario A, B, C: within-modality correlation, *r*_BPND_ = 0.638 and *r*_CBF_ = 0.481; Scenario D, E: *r*_BPND_ = 0.684 and *r*_CBF_ = 0.554.
- Cross-modality correlation: same regions correlation, Scenario A and C: *r*_region_ = 0.500; Scenario B and D: *r*_region_ = 0; Scenario E: *r*_region_ = 0.135; different regions correlation, Scenario A and C: *r*_id_ = 0.392 (leading to a conditional correlation of 0.250); Scenario B and D: *r*_id_ = 0; Scenario E: *r*_id_ = 0.113 (leading to a conditional correlation of 0.0574).

### Supplementary Material A3

Strategy 1 vs. Strategy 2 on a simulated dataset

To further illustrate the difference between Strategy 1 and 2 and how regional mean differences can affect Strategy 1, consider three datasets containing 24 participants each, simulated for Scenario A, B, and C, with the same starting value the random number generator (seed 12) corresponding to. The seed was chosen such that the estimated correlation with Strategy 2 approximately matched the marginal correlation defined by each scenario.

Supplementary Figure 1 displays the individual (first four columns) as well as the mean regional 5-HT2AR binding and CBF across all participants (fifth column). The correlation estimated by Strategy 1 is closely related to the regression slope shown in the fifth column. Visually, the data generated under Scenario B and C appear similar as most of the variation is driven by the large differences in regional mean, e.g., most patients have large BPND and CBF values in ventrolateral PFC and low values in caudate.

Supplementary Figure 2 displays the same data but normalized for each region, i.e., subtract the regional mean and divide by the regional standard deviation (across participants). This normalization does not have a large effect on Scenario A, but substantially affects Scenario B: the absence of correlation between BPND and CBF is much more obvious after removing regional effects.

Supplementary Figure 3 displays the same dataset, instead focusing on a subset of regions (the colored points in Supplementary Figure 1 match the points with numbers in Supplementary Figure 3). The regional difference in mean in Scenario B and C us apparent from the graphical display, yet does not affect the estimated Pearson correlation.

---

**Supplementary Figure 1.**
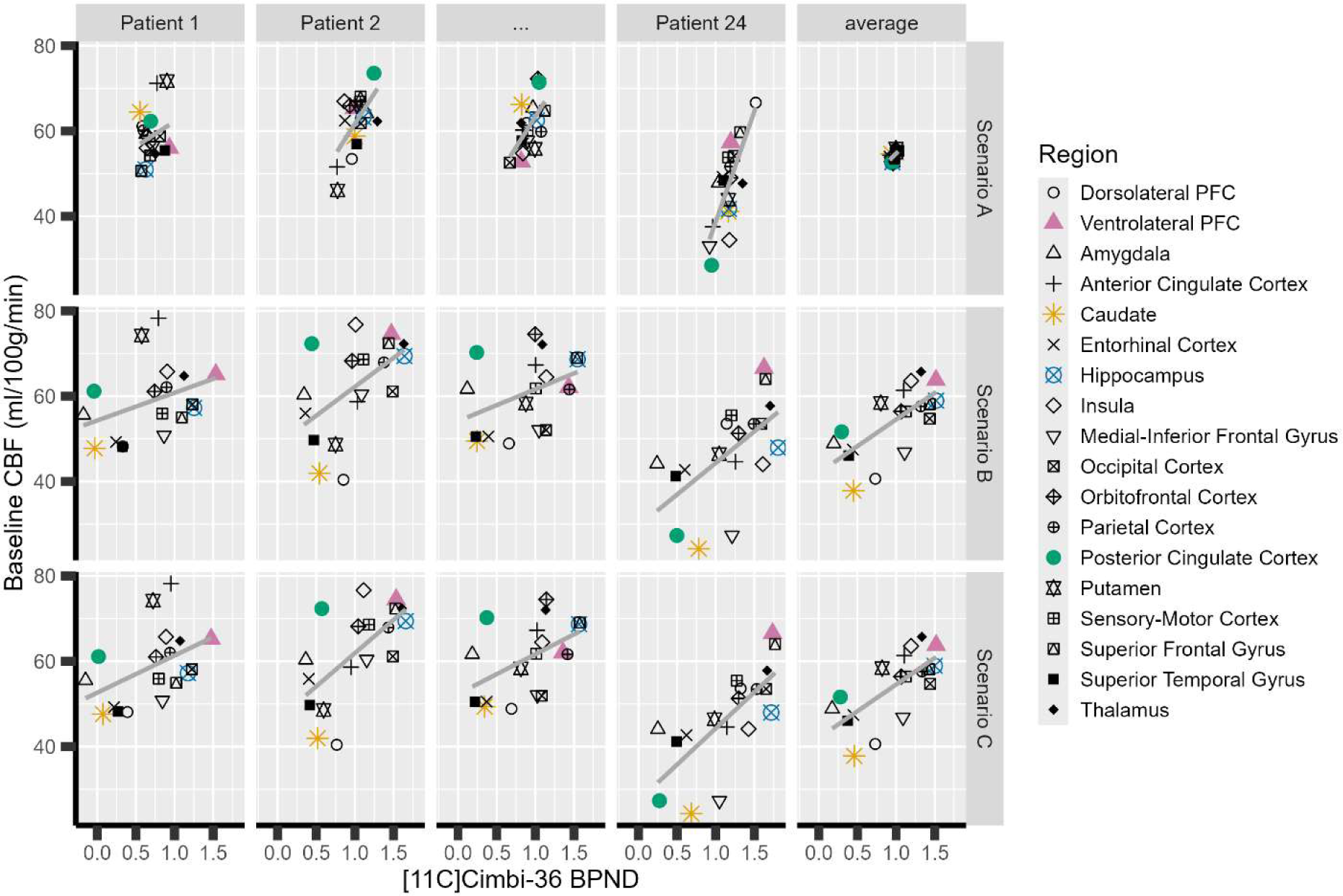
Scatterplot across regions of a simulated dataset for Scenario A, B, and C, with a different panel per participant (first four columns) or the average over participants (fifth column).

**Supplementary Figure 2.**
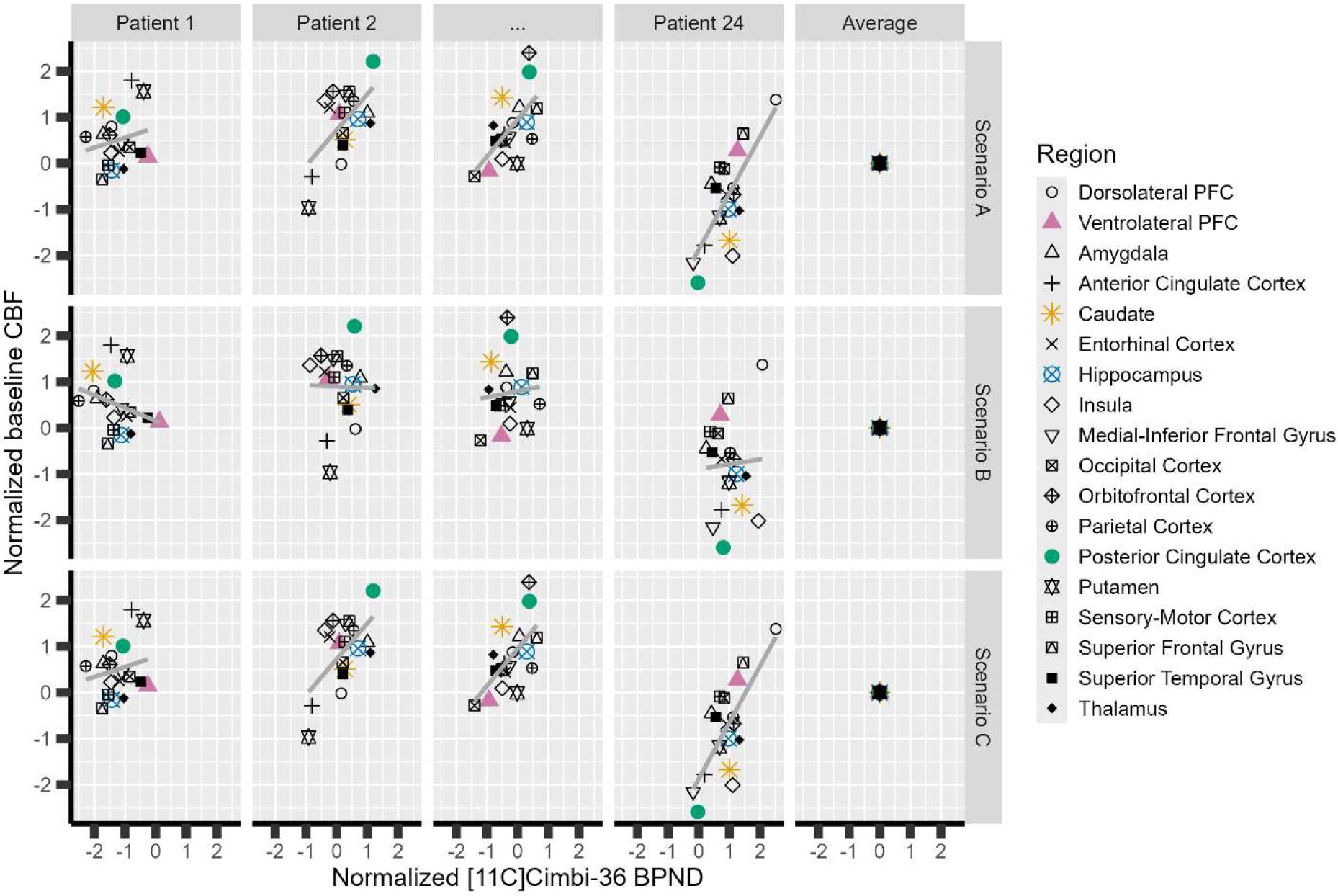
Same scatterplots as in Supplementary Figure 1, except that observations have been centered and scaled using region-specific means and standard deviations.

**Supplementary Figure 3.**
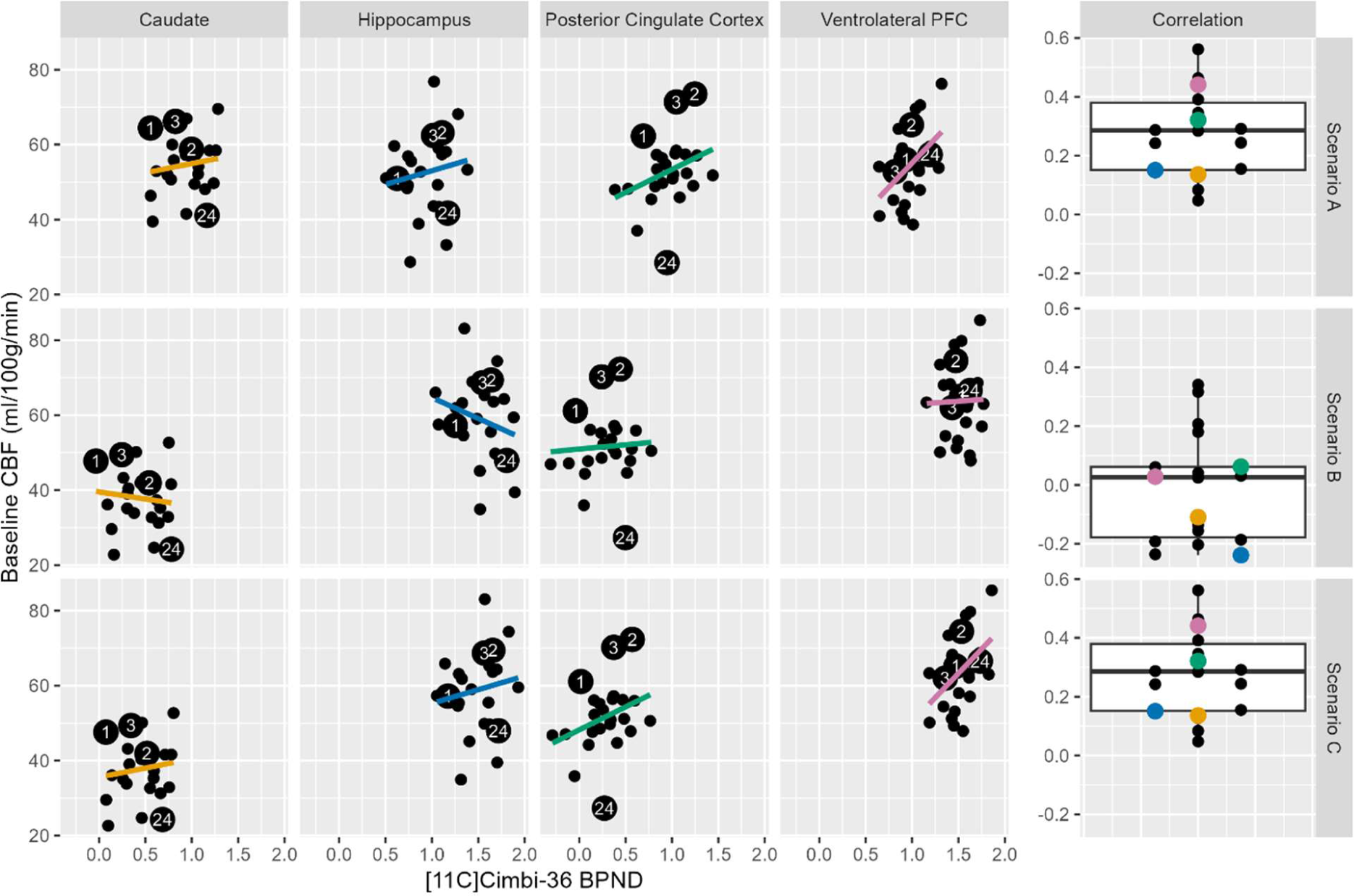
Scatterplot across participants of the same simulated dataset as in Supplementary Figure 1, except here with a different panel per region. The boxplot (rightmost column) displays the region-specific correlations which are closely related to the slopes shown in the scatterplots.

